# Aβ-42 sidechain deamidation at Q15 or N27 modulates protein aggregation and alters microglial cytokines and CD68

**DOI:** 10.1101/2025.10.22.684043

**Authors:** Martin N. Griffin, Daniel M. Dinakarapandian, Chenxing Li, Pranay Ramaswamy, Sia J. Saheba, Janie Lee, Tarunya Rao Sudarshan, Maria Sajimon, Christopher J. Wheeler, Jevgenij A. Raskatov, Anant K. Paravastu, Levi B. Wood

**Author notes:** Equally contributing authors (Martin N. Griffin, Daniel M. Dinakarapandian).

## Abstract

The progressive aggregation of amyloid beta (Aβ) monomers into oligomers is a critical factor in Alzheimer’s disease (AD) pathogenesis. Although mutated forms of Aβ have been shown to display altered aggregation dynamics, the specific effects of deamidated Aβ on microglial function remain understudied. Our research group previously found that the deamidated variant Aβ-42-N27D modified Aβ aggregation, reduced neurotoxicity, and reduced microglial reactivity, but the impact of Aβ-42 side chain deamidation in general on such parameters remained unclear. Here, we expanded on our prior work by investigating how two site-specific Aβ-42 mutations (Q15E & N27D), where neutral amide side chains are replaced with negatively charged carboxylic acids, affect aggregation and microglial immune response using a mouse microglial cell line. Size exclusion chromatography revealed that Aβ-42-Q15E and Aβ-42-N27D exhibit distinct aggregation profiles compared to Aβ-42 wild type (WT). Multiplexed analysis of 8 cytokines secreted into the culture medium revealed that Aβ-42-Q15E and Aβ-42-N27D decrease the expression of inflammatory cytokines such as IL-6, IP-10, and MIP-1α relative to Aβ-42-WT. Immunocytochemistry revealed that Aβ-42-Q15E and Aβ-42-N27D decrease CD68 expression relative to Aβ-42-WT. These findings demonstrate that deamidation significantly alters Aβ-42 aggregation and microglial activation, suggesting structural modifications to Aβ-42 modulate inflammatory signaling in AD. This work provides a foundation for future studies on Aβ-42 post-translational modifications as potential therapeutic targets in AD.S

## Introduction

Alzheimer’s disease (AD) is a progressive neurodegenerative disorder that affects millions of people each year and is the leading cause of dementia globally [1]. Accumulation of Aβ peptides into extracellular plaques is a defining hallmark of AD pathology [2]. While it remains unclear whether Aβ plaques directly drive pathology, Aβ oligomers as well as chronic neuroinflammation have both been strongly implicated in AD pathogenesis [3, 4]. Microglia are the resident immune cells of the brain and function to clear cellular debris, prune synapses, and maintain homeostasis under healthy conditions [5]. While microglia are believed to provide early protection during AD, microglial function becomes dysregulated during disease progression, reducing their ability to clear Aβ aggregates and contributing to a state of chronic neuroinflammation linked to cognitive decline [6, 7].

Given that Aβ aggregation state influences microglial activity [8], understanding how specific structural modifications to Aβ impact microglial activation is critical for elucidating mechanisms of inflammation in AD. In the healthy brain, microglia secrete neurotrophic factors that support neuronal health [9]. However, in AD, oligomeric forms of Aβ deregulate signaling pathways vital for cellular function and gene expression [10]. This dysregulation exacerbates chronic inflammation and accelerates neurodegeneration in AD patients [11]. Aβ interacts with microglial surface receptors, such as TREM2, CD36, and toll-like receptors, resulting in dysregulated phospho-protein signaling cascades that drive expression of inflammatory cytokines, including IL-1β, IL-6, and TNF-α [12].

In this study, we chose to focus on Aβ-42 as this form of Aβ is known to form more oligomers and be more neurotoxic than Aβ-40 [13]. Although limited, previous studies have explored how site-specific modification to Aβ variants influence aggregation and the resulting cellular responses [14]. Our recent study examined Aβ-42 with a substitution at position 27, where asparagine was replaced with aspartic acid (Aβ-42-N27D) and found that this variant exhibited reduced aggregation compared to wild-type Aβ-42, and notably produced substantially lower quantities of soluble oligomers [15]. In the current study, this result was successfully reproduced within a distinct laboratory environment, further validating the impact of Aβ-42-N27D on Aβ aggregation behavior. Our previous study also reported changes in microglial cytokine expression in response to the N27D variant. However, aspects of microglial activation and phagocytic function remained uncharacterized, providing the basis for our investigation into how deamidated Aβ-42 affects aggregation and broader microglial response.

In this study, we investigated how site-specific deamidation of Aβ-42 influences its aggregation kinetics and microglial immune response. We found that Q15E and N27D mutants of Aβ-42 exhibited reduced aggregation, lower expression of inflammatory cytokines, and diminished CD68 expression compared to Aβ-42-WT. Our data indicate that Aβ-42 deamidation at either N27 or Q15 modifies its aggregation and immune activation, highlighting deamidation as a potentially general mechanism for attenuating Aβ-induced inflammation.

## Results

### Deamidation of Aβ-42 at Q15 and N27 Alters Aggregation

We began the current study by investigating how site-specific deamidation affects Aβ-42 aggregation. To do so, we used size exclusion chromatography (SEC) to monitor the aggregation profiles of Aβ-42-Q15E, Aβ-42-N27D, and Aβ-42-WT over 24 hours. Freshly prepared monomers of each variant were isolated using SEC (**Fig. 1A**), diluted to 100 μM, and incubated at 37°C for 24 hours to allow aggregation to occur. The resulting aggregation profiles were then analyzed by re-injection onto the SEC column.

**Figure 1.**
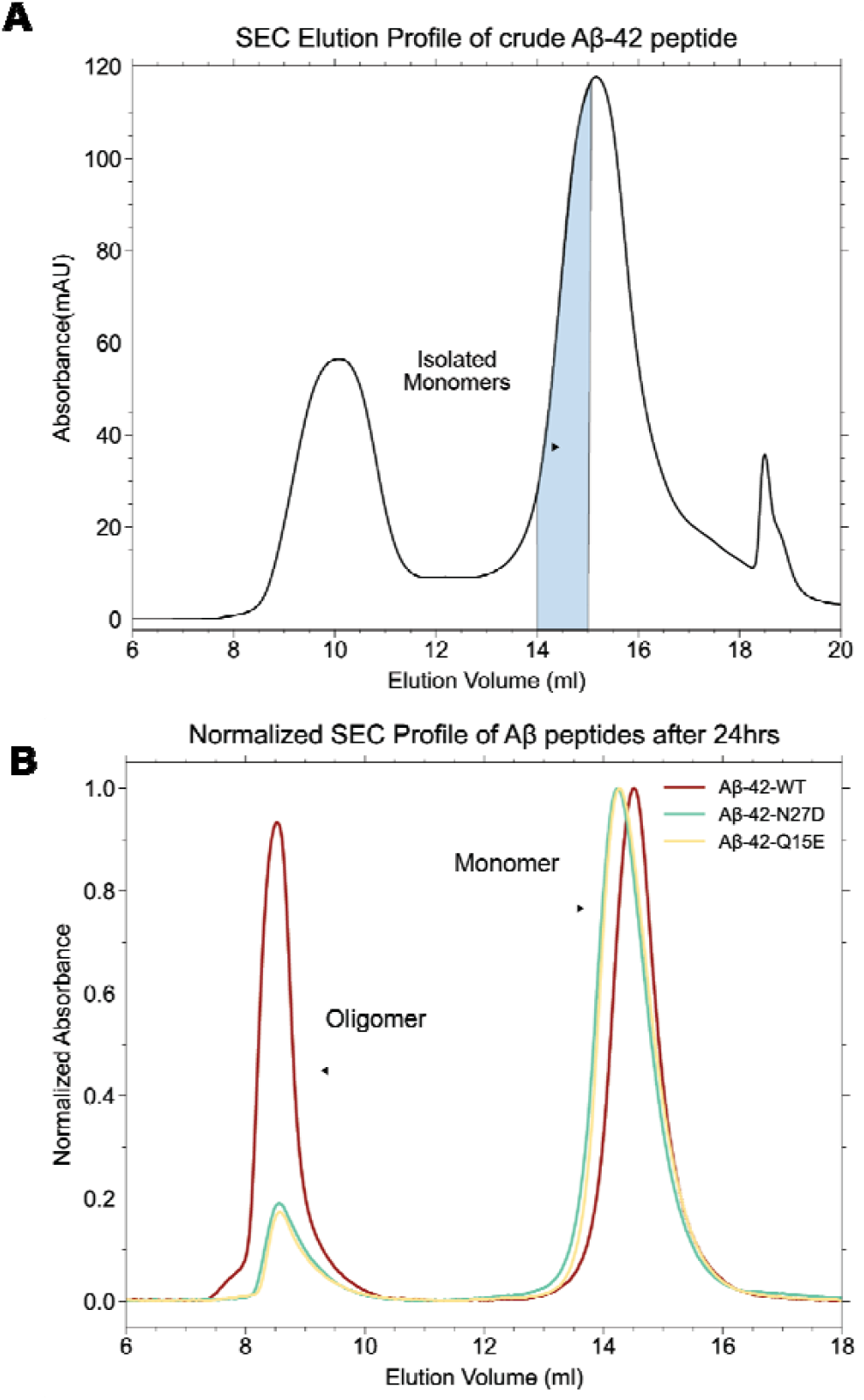
Deamidation of Aβ-42 at Q15 and N27 Leads to Less Oligomer Formation. **A**. Aβ-42 monomers isolation using Size Exclusion Chromatography. Crude Aβ-42 peptides wer dissolved in 50mM NaOH and injected into SEC column at 0.5 mL/min flow rate. The monomers eluted at 14-15 mL (highlighted area in blue) were collected and diluted to the final concentration of 100 μM. The collected monomers were incubated at 37°C for 24 hours. **B**. SEC profiles showing difference in aggregates formed from incubated monomers of Aβ-42 WT, Aβ-42 N27D, and Aβ-42 Q15E after 24 hours.

The SEC elution profiles revealed distinct aggregation behaviors among the three variants (**Fig. 1B, Sup Fig. 2**). Aβ-42-WT displayed a pronounced oligomeric peak at approximately 9 ml elution volume, indicating substantial formation of higher-order aggregates after 24 hours of incubation. The deamidated variants, Aβ-42-N27D and Aβ-42-Q15E, showed reduced oligomeric peak formation, with peak heights approximately 75% lower than the wild-type variant. The monomeric peaks (eluting at ∼14 - 15 ml) remained prominent for both mutants, suggesting that deamidation inhibits the progression from monomeric to oligomeric states.

**Figure 2.**
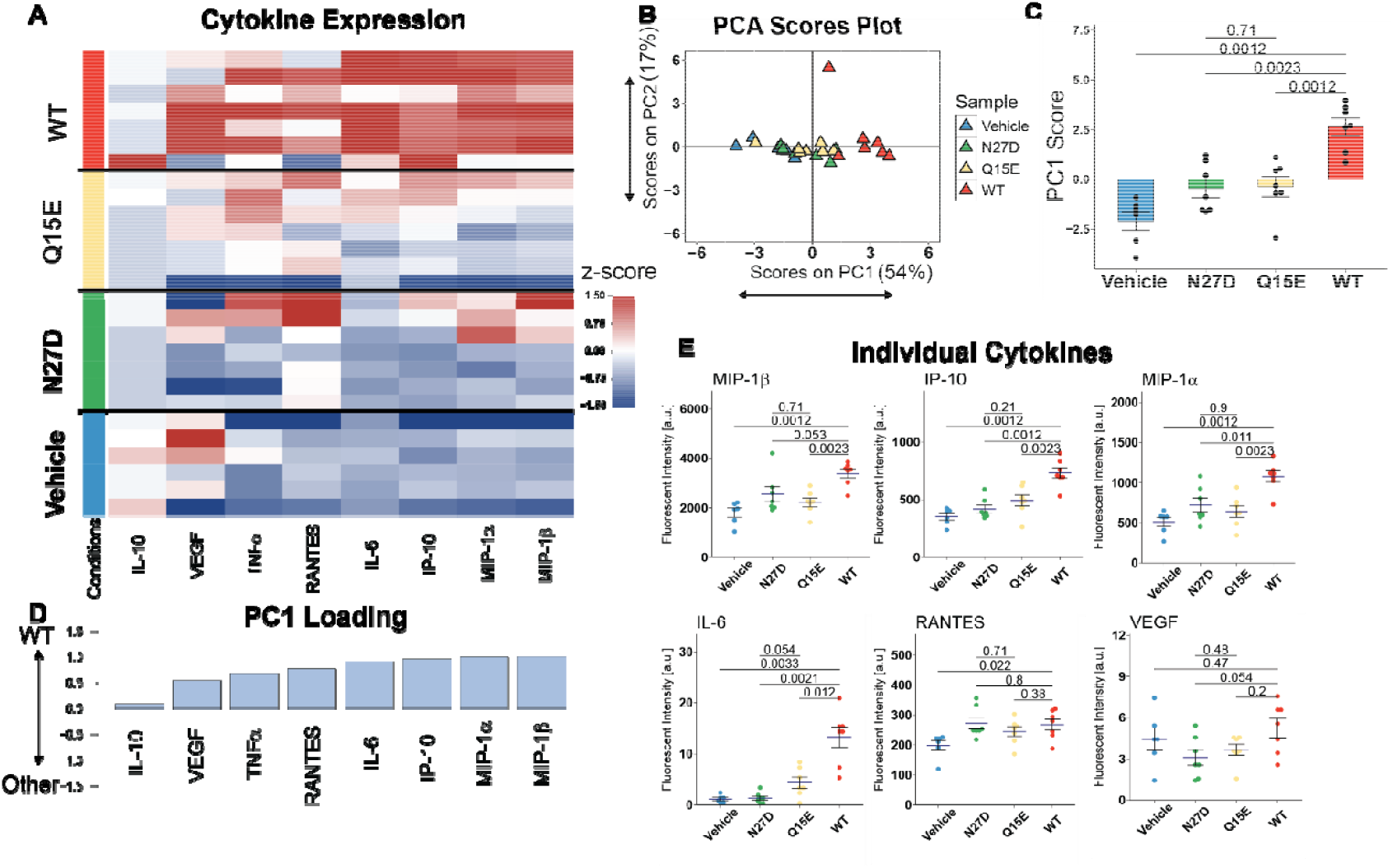
Deamidated Aβ modulates cytokine expression in mouse microglial cell line. **A**. Multiplexed Luminex analysis of 8 cytokines (columns) expressed into the culture medium after 24 hours of incubation with 50nM of each Aβ variant (n=6-8 wells/group). Each row in the z-scored heat map denotes an individual culture sample well. **B**. Scores plot of principle component analysis (PCA). Each triangle represents a sample well. PC1 explained 54% of the variance, and PC2 explained an additional 17%. **C**. Bar plot of PC1 scores for each group. In all bar plots, dots denote individual samples. **D**. PCA identified a variable, PC1, that captured the greatest axis of variation across experimental groups. PC1 consisted of a profile of cytokine that were upregulated with Aβ-42-WT. **E**. Dot plots for individual cytokines (mean ± SEM Wilcoxon rank sum test). In all barplots, dots denote individual samples. Each dot denotes an individual sample well. All p-values reflect Wilcoxon rank sum tests with Bonferroni adjustment for multiple comparisons. a.u., arbitrary units.

### Deamidation of Aβ-42 modulates cytokine expression in mouse microglial cell line

Chronic microglial activation is a hallmark of AD pathology and is closely linked to Aβ oligomerization [16]. We therefore next investigated whether deamidated Aβ-42 variants would elicit distinct cytokine responses compared to Aβ-42-WT. To do so, we applied a physiologically relevant concentration (50nM [17, 18]) of Aβ-42-WT, Aβ-42-N27D, and Aβ-42-Q15E to mouse microglial cells (MMCs) for 24 hours. We then quantified 8 inflammatory cytokines associated with chronic inflammation secreted into the culture medium using a Luminex multiplexed immunoassay [19, 20] (**Fig. 2A**).

Principal component analysis (PCA) identified principal components (PC1 and PC2) that captured the greatest variance in cytokine expression across samples, revealing clear separation between deamidated variants and Aβ-42-WT (**Fig. 2B**). Scores on PC1 showed that Aβ-42-WT clustered separately on the far right, both Aβ-42-N27D and Aβ-42-Q15E clustered in the middle, while Vehicle clustered on the left. Similarly, PC1 scores showed significant upregulation in Aβ-42-WT compared to Vehicle, Aβ-42-N27D, and Aβ-42-Q15E (**Fig. 2C**), indicating the effect of Aβ-42-WT was normalized with Aβ deamidation. PC1 consisted of a cytokine profile and the loadings on PC1 reflected the relative influence of each cytokine. MIP-1β, MIP-1α and IP-10 had the highest absolute loadings on PC1, indicating their most significant contribution to the PC1 score. Among the top contributors to PC1 (**Fig. 2D**), MIP-1β, MIP-1α, IP-10, and IL-6 were significantly upregulated in Aβ-42-WT compared to vehicle, Aβ-42-N27D, and Aβ-42-Q15E **(Fig. 2E, Sup Fig. 3**). Together, both Aβ-42-N27D and Aβ-42-Q15E elicited lower cytokine expression compared to Aβ-42-WT, suggesting that Aβ deamidation may mitigate microglial dysregulation in AD.

**Figure 3.**
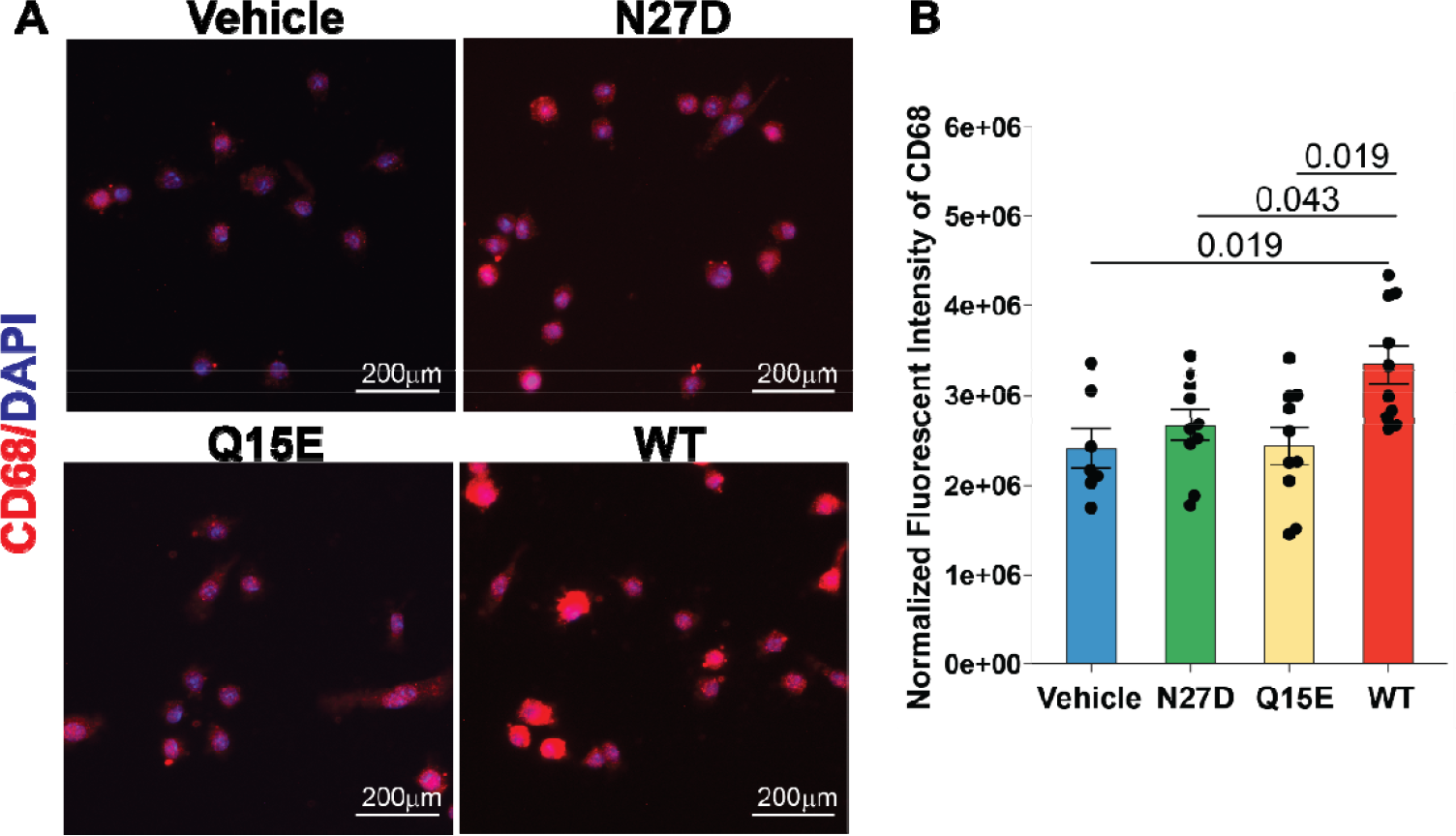
Deamidation abrogates Aβ-induced CD68 expression in mouse microglial cell line. **A**. ICC for CD68 (red) and DAPI (blue) after 2 hours of incubation with 50nM of each Aβ variant (scale bar: 200 μm, representative sections from n=12-13 sample well/group). **B**. Integrated CD68 florescent intensity normalized by number of nuclei (n=12-13, mean±SEM, Wilcoxon rank sum tests with Bonferroni adjustment for multiple comparisons).

### Aβ-42 deamidation attenuates CD68 expression in mouse microglial cell line

Aβ-42 oligomers have been shown to increase microglial activation and dysregulate phagocytosis [21]. CD68, a lysosomal marker, is consistently shown to be upregulated in activated microglia [22]. Therefore, we utilized immunocytochemistry (ICC) to stain for CD68 in MMCs treated with 50nM of Aβ-42-WT, Aβ-42-N27D, and Aβ-42-Q15E for 2 hours, as this is a time period where studies have shown increased levels of Aβ-42 uptake [23]. ICC examination of CD68 showed that CD68 labeling is reduced in the Vehicle, Aβ-42-N27D, and Aβ-42-Q15E (**Fig. 3A, Sup Fig. 4**). Quantification of CD68 fluorescent intensity normalized to DAPI cell count revealed reduced CD68 expression in Aβ-42-N27D and Aβ-42-Q15E compared to Aβ-42-WT (**Fig. 3B**), indicating that Aβ deamidation reduces its potency to induce microglial activity.

## Discussion

Aβ oligomers are widely recognized as drivers of microglial activation, inflammation, and neurodegeneration [16, 24]. Thus, Aβ mutants that promote oligomerization would be expected to induce stronger microglial activation than mutants that favor monomeric forms. Building on our previous study where we examined the impact of Aβ-42-N27D on aggregation and microglial activity [15], this work differs methodologically in that we freshly prepared our samples by isolating monomers using SEC to remove any pre-seeding effects that could affect the aggregation kinetics and behavior of the Aβ-42 peptide. This modification led to a more consistent microglial response than observed previously.

Our SEC data indicates that Aβ-42-Q15E and Aβ-42-N27D aggregate less than Aβ-42-WT. Therefore, the observed differences in microglial activity could be partially due to altered oligomer formation. Similar findings have been reported for Aβ-40, where the Aβ-40-Q15E and Aβ-40-N27D variants showed slower aggregation compared to Aβ-40-WT [14]. Thus, if microglial activation were solely dependent on aggregation state, these findings would support the conclusion that Aβ-42-Q15E and Aβ-42-N27D are less activating than Aβ-42-WT.

While the aggregation dynamics of deamidated mutants of Aβ-40 have been documented [14], the aggregation dynamics and microglial response to Aβ-42-Q15E and Aβ-42-N27D remains poorly understood. Our SEC data shows that Aβ-42-Q15E aggregates similarly to Aβ-42-N27D, despite the Q15E switch occurring in a different region of Aβ-42 than N27D. Research has shown that the loss of negative charges at specific residues on Aβ-42 leads to increased aggregation, enhanced stability, and greater neurotoxicity of the peptide [25]. Given that both N27D and Q15E introduce a negative charge to Aβ-42, this supports the idea that deamidation at these residues could result in slower aggregation. A simple explanation is that WT is net trianionic, while N27D and Q15E are net tetra-anionic. The higher negative charge is expected to enhance water solubility and disfavor hydrophobic collapse that drives oligomer formation [26]. Another effect added on top is the loss of fibrillogenicity of the Q15 or N27 sidechain upon deamidation. Future structural work may reveal additional, more subtle contributing factors.

One major finding in our study is that both Aβ-42-Q15E and Aβ-42-N27D reduce microglial activation and diminish expression of inflammatory cytokines compared to Aβ-42-WT. As Aβ-42-WT forms more oligomers than Aβ-42-Q15E and Aβ-42-N27D, Aβ-42-WT could be able to bind to more microglial surface receptors than the deamidated mutants. The reduced oligomer populations in Aβ-42-Q15E and Aβ-42-N27D could explain the decreased inflammatory cytokine expression observed in the deamidated mutants [27, 28]. While limited differences were found between Aβ-42-Q15E and Aβ-42-N27D in terms of cytokine expression, our data suggest that Aβ-42-Q15E and Aβ-42-N27D may interact differently with microglia [29]. One such analyte, IL-6, is expressed significantly higher in microglia treated with Aβ-42-Q15E than Aβ-42-N27D. Since IL-6 production is regulated by multiple receptor-mediated signaling pathways [30, 31], these findings suggest that Aβ-42-Q15E and Aβ-42-N27D may engage distinct microglial surface receptors. This suggestion is plausible given that the deamidation sites differ between the two mutants, potentially altering their conformation and resultant receptor binding.

In addition to cytokine profiling, we quantified CD68 because it is upregulated in AD pathology and cognitive decline [17]. We found that both Aβ-42-Q15E and Aβ-42-N27D induce significantly lower expression of CD68 compared to Aβ-42-WT. Since CD68 is a marker of lysosomal activity, it also serves as an indirect measure for evaluating lysosomal degradation of Aβ-42 [22]. As the expression of CD68 is lower in the deamidated variants, this may indicate that Aβ-42-Q15E and Aβ-42-N27D are more efficiently degraded or result in reduced lysosomal activation compared to Aβ-42-WT.

The N27D and Q15E mutations were designed to mimic spontaneous deamidation events that disrupt aggregation. Both asparagine and glutamine residues contribute to fibril stability through interdigitated hydrogen bonding networks observed in fibril structures [32, 33]. The N27D mutation substitutes asparagine with aspartic acid at residue 27 within the turn region (residues 22-29), a structurally sensitive area important for β-hairpin formation during aggregation and oligomerization [32-36]. Similarly, Q15E introduces a negative charge within the central hydrophobic core, where Q15 normally contributes to fibril stability [14, 32]. The introduction of negatively charged residues at these positions likely disrupts aggregation through electrostatic repulsion and loss of stabilizing hydrogen bonding interactions, consistent with previous observations that charge manipulations at specific residues modulates Aβ aggregation propensity and neurotoxicity. Previous studies demonstrating that the loss of negative charges at specific residues leads to increased aggregation and enhanced neurotoxicity [37] support our observation that the addition of negative charges through deamidation has the opposite effect. Therefore, the reduced cytokine and CD68 expression observed in the microglia treated with either deamidated variant could be due to the addition of the negative charge. However, detailed structural characterization will be necessary to fully elucidate the mechanisms by which these deamidation-mimicking mutations alter Aβ-42 aggregation behavior.

While our study provides important insights into how deamidated Aβ-42 mutants modulate microglial activation, several limitations should be acknowledged. First, the experiments were conducted in an immortalized microglial monoculture, which shares key behavioral characteristics with primary microglia but does not fully capture the complexity of primary microglial phenotypes [38], particularly in the context of AD. Additionally, although CD68 staining modulation suggests deamidation altered phagocytic activity, CD68 primarily reflects lysosomal activity and does not directly measure microglial Aβ uptake via phagocytosis, a process heavily implicated in AD pathology [39]. Also, our cytokine profile suggests that the two deamidated Aβ-42 variants may elicit slightly different microglial responses, motivating future analysis of receptor binding kinetics or downstream signaling activation. Together, these limitations highlight the need for in vivo studies and expanded mechanistic analyses.

Overall, our data show that deamidation of Aβ-42 reduces aggregation and dampens microglial activation. Given the growing clinical significance of Aβ aggregation and immune activity in AD [40], our findings therefore support further interrogation of Aβ-42 deamidation as a clinically relevant approach to reducing its toxicity. In particular, the formulation of new pharmacological strategies for post-translational deamidation could represent a viable strategy for AD treatment.

## Methods

### Aβ Preparation

Aβ-42-WT, Aβ-42-N27D, and Aβ-42-Q15E peptides were synthesized using a CEM Liberty Blue peptide synthesizer in-house following Fmoc solid-phase peptide synthesis protocol. Following synthesis, crude peptides were verified for molecular weight using electrospray ionization mass spectrometry (ESI-MS) at the proteomics facility, confirming the presence of peptides with correct molecular masses corresponding to their expected sequences. See **Supporting Information Figure 1**.

### Monomer Isolation

For monomer isolation, lyophilized peptides were dissolved in 50 mM sodium hydroxide (NaOH) and allowed to equilibrate for 20 minutes at room temperature to ensure complete dissolution. The peptide solutions were then immediately subjected to size exclusion chromatography (SEC) to isolate monomeric fractions.

### Size Exclusion Chromatography

Size exclusion chromatography (SEC) was performed using a Superdex 75 increase 10/300 GL column (GE Healthcare) connected to an ÄKTA GO chromatography system. The running buffer consisted of 20mM sodium phosphate (NaPi, pH 7.4). Peptide samples (3 mL) were injected at a flow rate of 0.5 mL/min, and elution was monitored by UV absorbance at 280 nm. Monomeric fractions were identified based on their elution volume (∼14 mL) and collected accordingly. The isolated monomers were immediately diluted to a final concentration of 100 μM in the same buffer and used for aggregation studies.

For aggregation analysis, freshly isolated monomers were incubated at 37°C for 24 hours without agitation. Following incubation, samples were re-injected into the SEC column under identical conditions to analyze the aggregation state.

### Cell Conditioning and Media Collection

Mouse microglial cells (MMCs) were cultured in DMEM-F12 medium supplemented with 10% fetal bovine serum and 1% Penicillin-Streptomycin until they reached 70% confluency. Cells were stored in an incubator with 95% air and 5% CO2 atmosphere at 37□°C. Upon reaching 70% confluency, cells were seeded into 12-well plates and incubated for 24 hours. Cells were treated with either 50 nM Aβ42-WT, 50 nM Aβ42-N27D, 50nM Aβ42-Q15E or vehicle (20mM NaPi) for 24h followed by medium collection. Collected medium was centrifuged at 13,000 RPM for 10 min at 4°C, then the supernatant was collected and stored at-80°C. Multiplexed cytokine quantification was conducted using the Milliplex® MAP Mouse Cytokine/Chemokine 18-Plex kit (Eotaxin^*^, GM-CSF^*^, IFN-γ^*^, IL-1β^*^, IL-2^*^, IL-3^*^, IL-4^*^, IL-6, IL-10, IL-12p70^*^, IL-13^*^, IP-10, M-CSF^*^, MIP-1α, MIP-1β, RANTES, TNFα, and VEGF) (Millipore Sigma MCYTOMAG-70K). Cytokines marked with an asterisk did not fall within a linear range of bead fluorescent intensity vs. protein concentration and were not included in the analysis.

### Multivariate Analysis of Cytokine Data

Data were analyzed and figures were generated via RStudio (Boston, MA) using the R programming language. The tidyverse collection of packages was used to conduct data processing. R package heatmap3 were used to generate heatmaps; R packages ggplot2, ggpubr, ggbeeswarm, and ggprism were used to generate dotplots and barplots. Clustering was conducted using the hclust function of the stats package in R using Euclidean distance in the unweighted pair group method with arithmetic mean. For functional and protein analysis, Bonferroni adjustment for applied to correct for multiple comparisons.

Multivariate cytokine data were analyzed using Principal Component Analysis (PCA) to identify axes, called principal components (PCs), that capture the greatest variance in cytokine expression across samples. PCA was performed using the R package ropls, with all data z-scored prior to analysis. Two principal components were extracted, and 5-fold cross-validation was applied to evaluate model robustness. All cytokine data were z-scored prior to analysis. PCA loadings were used to represent the contribution of each analyte to the observed variance.

### Cell Conditioning for CD68 Analysis

Mouse microglial cells were cultured in DMEM-F12 medium supplemented with 10% fetal bovine serum and 1% Penicillin-Streptomycin until they reached 70% confluency. Cells were stored in an incubator with 95% air and 5% CO2 atmosphere at 37 °C. Upon reaching 70% confluency, cells were seeded into a 96-well plate and incubated for 24 hours. Cells were treated with either 50 nM Aβ42-WT, 50 nM Aβ42-N27D, 50nM Aβ42-Q15E or vehicle (20mM NaPi) for 2h followed by fixing with 4% PFA on ice for 10 minutes. Cells were then rinsed 3x with cold PBS and then stored at 4°C.

### Immunocytochemistry

Fixed cells were permeabilized with 0.01% Triton X-100 in PBS for 10 minutes at room temperature, then rinsed twice with PBS. Cells were blocked for 1 hour at room temperature using blocking buffer containing 5% BSA and 3% goat serum in PBS. Following blocking, cells were incubated overnight at 4°C with rat anti-CD68 antibody (1:200; ThermoFisher). The following day, cells were washed five times with PBS containing 5% BSA, with each wash lasting 5 minutes. Wells were then incubated with Alexa Fluor 555 conjugated secondary antibody (1:2000; ThermoFisher) for 2 hours at room temperature. Afterward, cells were washed three times with 5% BSA in PBS for 5 minutes each wash. Nuclei were counterstained with DAPI for 20 minutes at room temperature, followed by three additional 5-minute washes with 5% BSA in PBS. Cells were then rinsed three times with PBS and stored in PBS until imaging. Samples were imaged using epifluorescent microscopy on a Zeiss Axio Observer Z.1 inverted microscope with a 20x lens and halogen bulb illumination using Zeiss filter set 49 to image DAPI and Zeiss filter set 20 to image Alexa fluor 555. Quantification of CD68 florescent intensity was analyzed by dividing the CD68 intensity by DAPI cell counts (ImageJ Macro).

### ImageJ Macro

An ImageJ macro was developed to automate the quantification of CD68 fluorescent intensity. The macro recursively scans a directory for merged images containing targeted red (CD68) and blue (DAPI) channels, while excluding single-channel files. Each image was then split into individual channels. The DAPI channel is thresholded using the Otsu method and segmented via watershed to count DAPI-positive nuclei, while measuring the red channel integrated fluorescent intensity. The intensity value was then normalized to the number of DAPI-positive nuclei to account for cell density. Finally, output values were printed to a log for export.

## Data Availability

The data described in the manuscript are located within the manuscript and supporting information.

### Supporting Information

This article contains supporting information.

