## Supplemental Information for "Aβ-42 sidechain deamidation at Q15 or N27 modulates protein aggregation and alters microglial cytokines and CD68"

+ Equally contributing authors

* Corresponding Authors

**Supporting Information**

**Supporting Information Tables**

**Supporting Table 1. Antibody Information**

| Antibody | Supplier | Catalog # | Dilution |
| --- | --- | --- | --- |
| CD68 | ThermoFisher | 14-0681-82 | 1:200 |

**Supporting Information Figures**

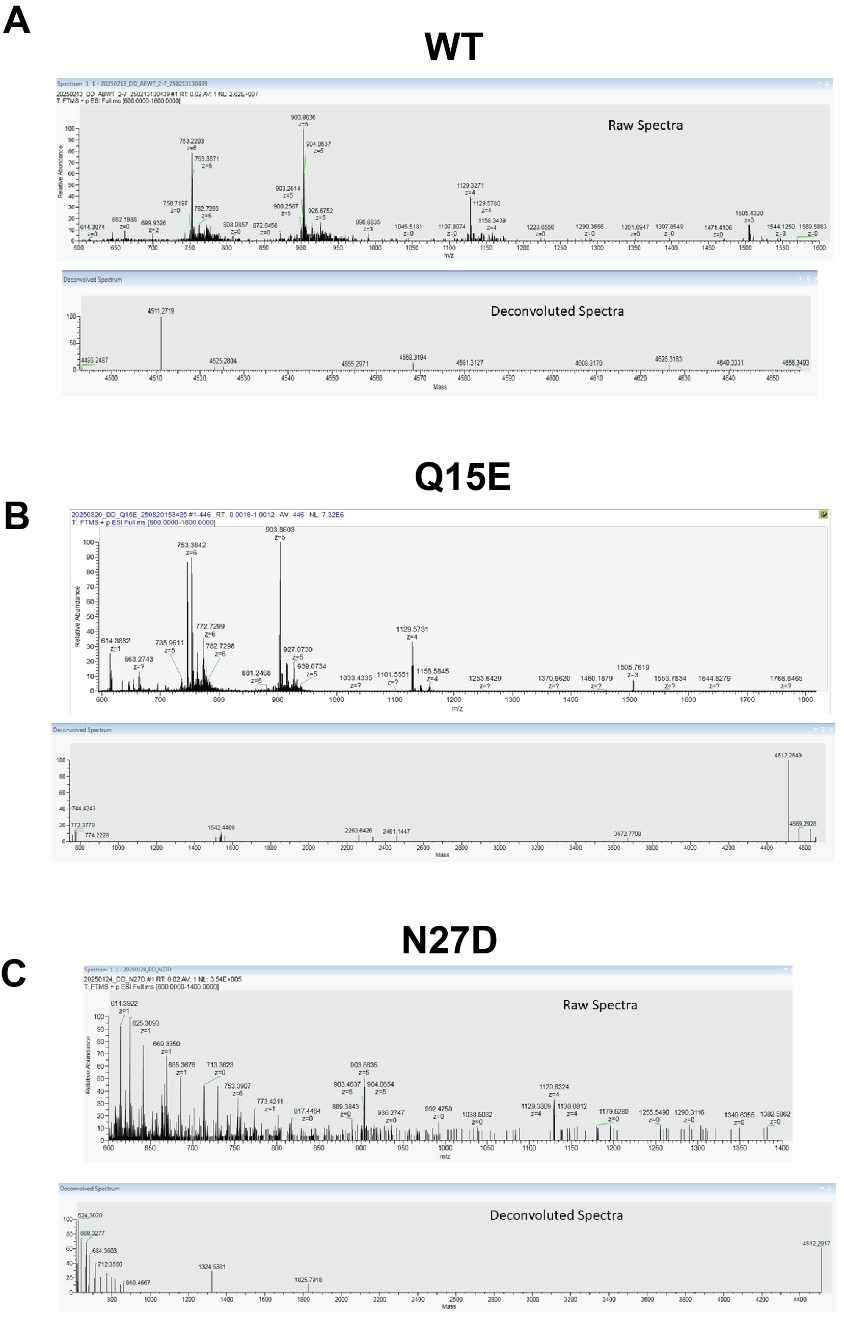

**Supporting Figure 1. ESI-MS analysis of synthetic peptides. A.** Raw and Deconvoluted Mass spectra of Aβ-42-WT with a monoisotopic mass of 4511.27 Da. **B.** Raw and Deconvoluted Mass spectra of Aβ-42-Q15E with a monoisotopic mass of 4512.26 Da. **C.** Raw and Deconvoluted Mass spectra of Aβ-42-N27D with a monoisotopic mass of 4512.26 Da.

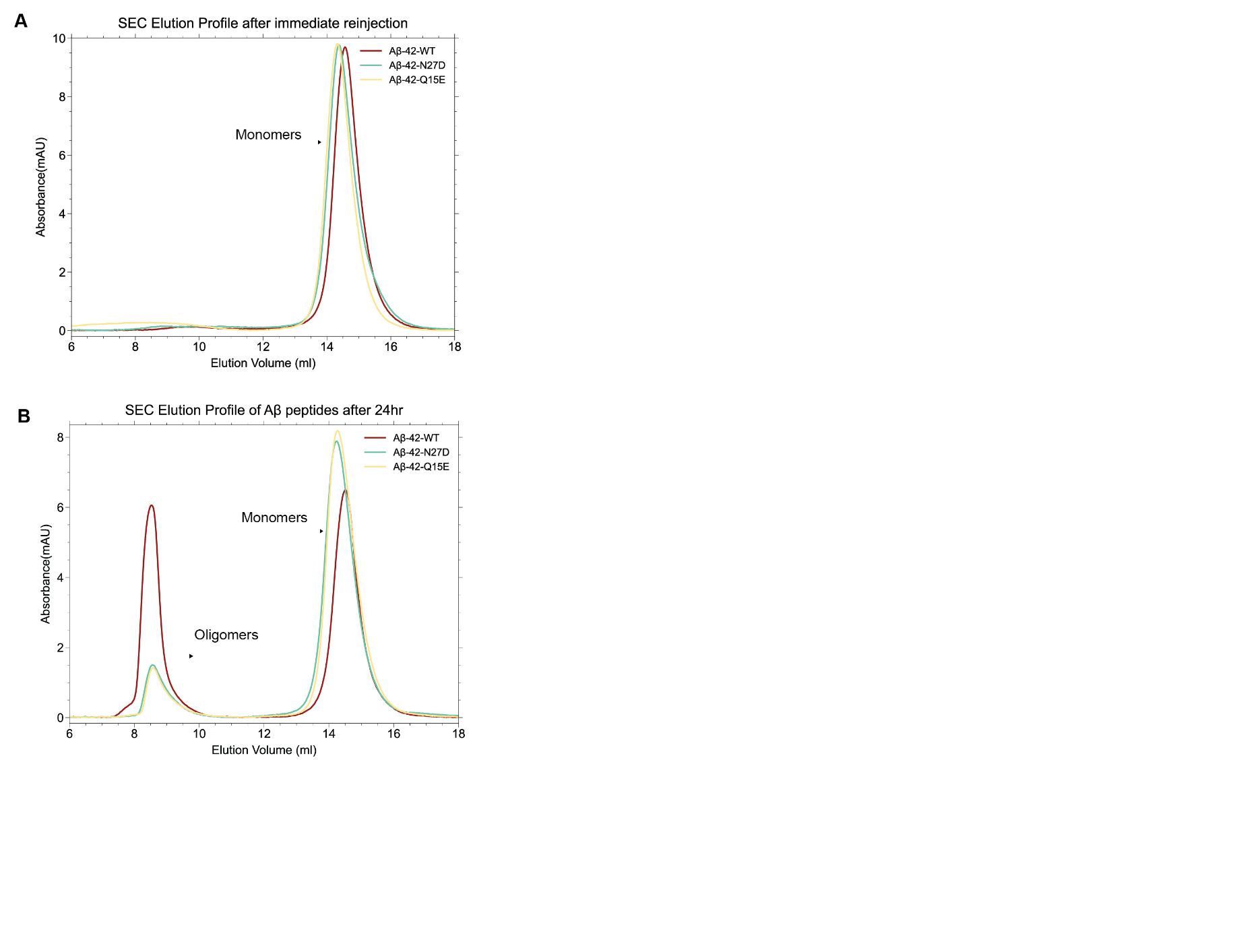

**Supporting Figure 2. Non-normalized SEC profiles of Aβ-42 peptides. A.** Immediate reinjection of freshly isolated monomers at t = 0 hr. **B.** SEC profiles of incubated monomers of Aβ-42-WT, Aβ-42-Q15E, and Aβ-42-N27D after 24 hours at 37°C.

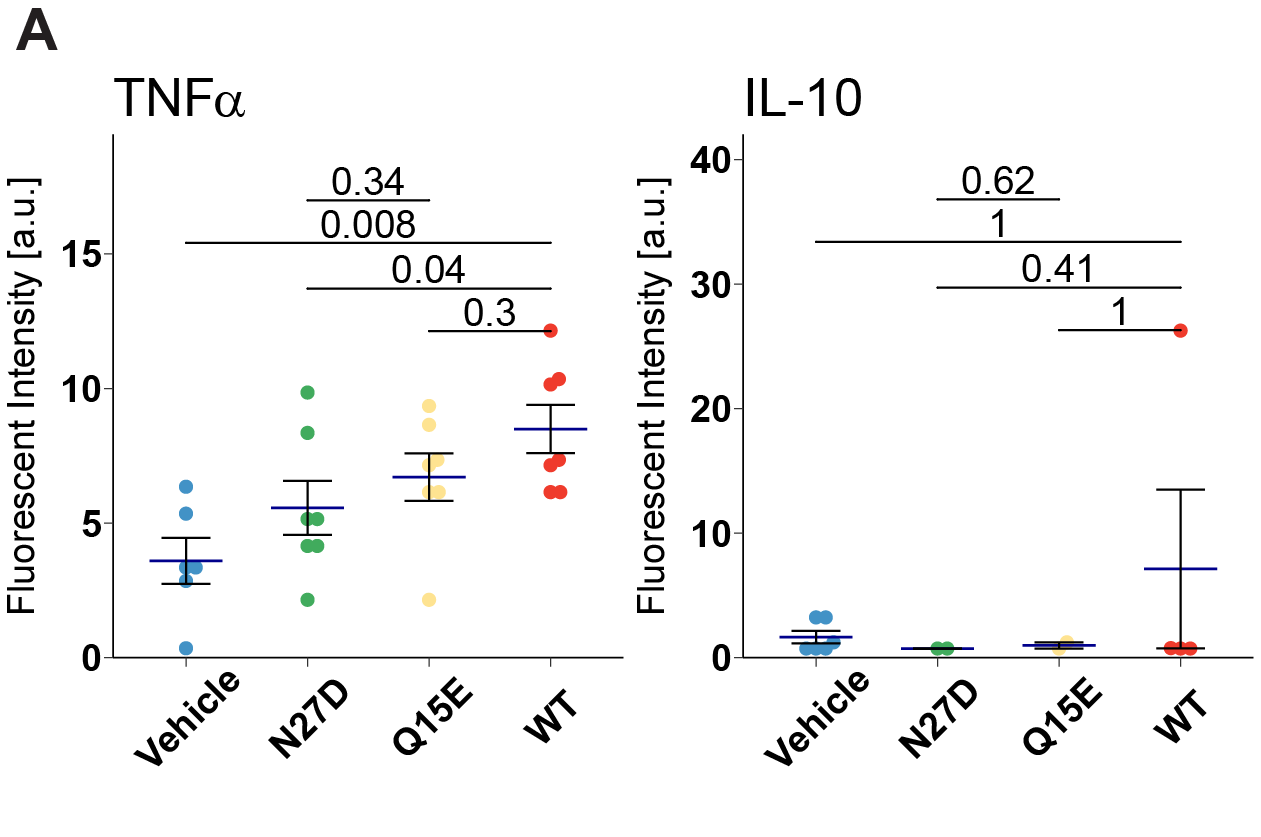

**Supporting Figure 3. Deamidated Aβ modulates cytokine expression in mouse microglial cell line. A.** Dot plots for individual cytokines (mean ± SEM Wilcoxon rank sum test). In all barplots, dots denote individual samples. Each dot denotes an individual sample well. mean±SEM. All p-values reflect Wilcoxon rank sum tests with Bonferroni adjustment for multiple comparisons. a.u., arbitrary units.

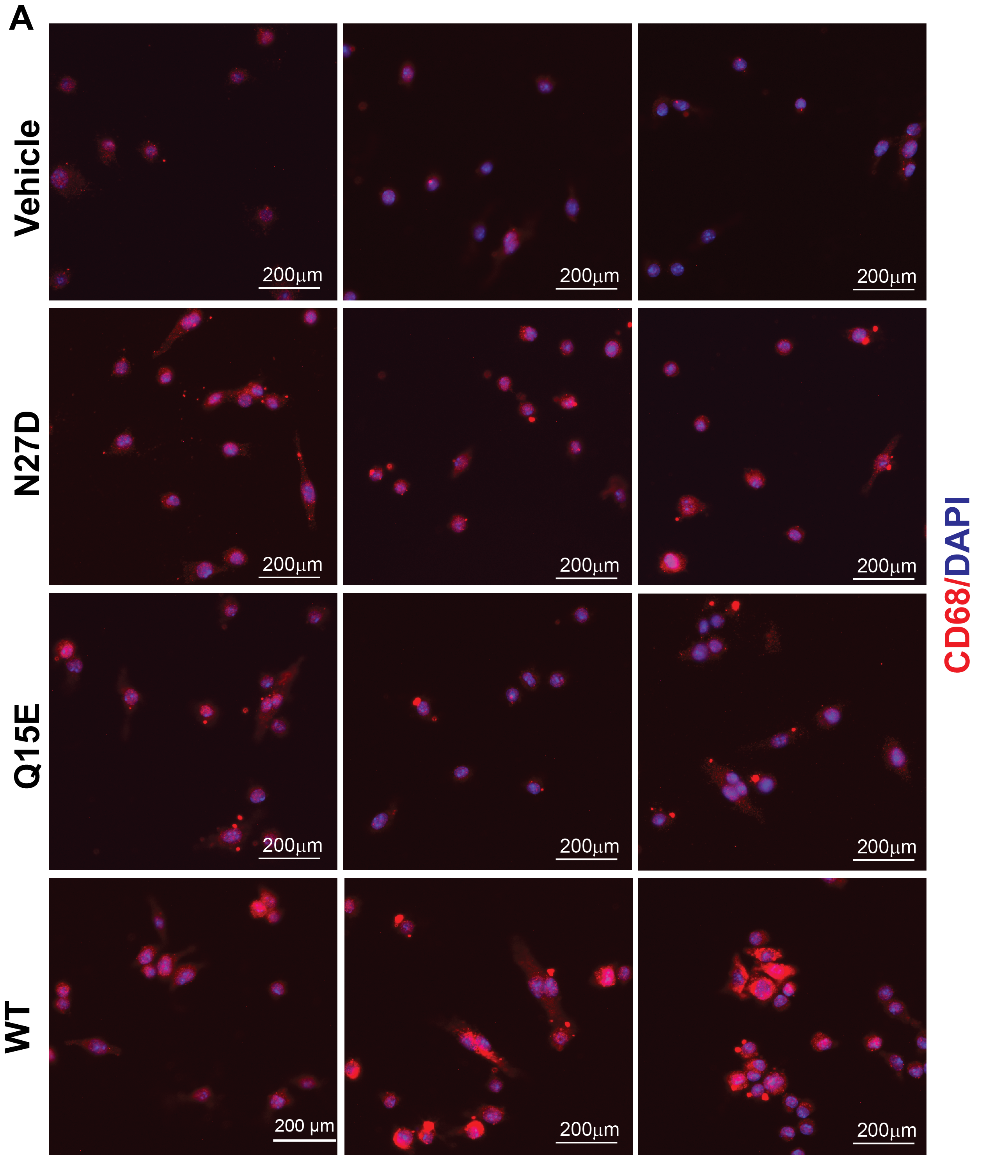

**Supporting Figure 4. Deamidation abrogates Aβ-induced CD68 expression in mouse microglial cell line. A**. Representative ICC for CD68 (red) and DAPI (blue) after 2 hours of incubation with 50nM of each Aβ variant (scale bar: 200 μm, representative sections from n=12-13 sample well/group).
